# Flies improve the salience of iridescent sexual signals by orienting toward the sun

**DOI:** 10.1101/2020.05.09.085639

**Authors:** Thomas E. White, Tanya Latty

## Abstract

Sunlight is the ultimate source of most visual signals. Theory predicts strong selection for its effective use during communication, with functional links between signal designs and display behaviours a likely result. This is particularly true for iridescent structural colours, whose moment-to-moment appearance bears a heightened sensitivity to the position of signallers, receivers, and the sun. Here we experimentally tested this prediction using *Lispe cana*, a muscid fly in which males present their structurally coloured faces and wings to females during ground-based sexual displays. In field-based assays we found that males actively bias the orientation of their displays toward the solar azimuth under conditions of full sunlight and do so across the entire day. This bias breaks down, however, when the sun is naturally concealed by heavy cloud or experimentally obscured. Our modelling of the appearance of male signals revealed clear benefits for the salience of male ornaments, with a roughly four-fold increase in subjective luminance achievable through accurate display orientation. These findings offer fine-scale, causal evidence for the active control of sexual displays to enhance the appearance of iridescent signals. More broadly, they speak to predicted coevolution between dynamic signal designs and presentation behaviours, and support arguments for a richer appreciation of the fluidity of visual communication.

## Introduction

The structure of any effective communication system is shaped its response to basic challenges. Encoding information (Weaver et al. 2018; White 2020), ensuring its delivery (Endler and Thery 1996), and separating signal from noise (Warrant 2016) are essential features of signalling systems, and selection has generated diverse solutions. In general terms, theory predicts the coevolution of signals and signalling behaviours that secure the exchange of information between interested parties while minimising exposure to eavesdroppers (Endler 1992; Endler 1993). Males of the common eggfly *Hypolimnas bolina*, for example, combine limited-view iridescent colours with precision display behaviours to deliver maximally conspicuous signals during courtship (White et al. 2015). Great bowerbirds, by contrast, present coloured objects to females from within constructed viewing ‘theatres’, and in doing so control both the object of display and the broader visual environment (Endler et al. 2010; Kelley and Endler 2012).

Sunlight is the ultimate source of most visual signals, as its colour and intensity shape the appearance of reflective surfaces. Selection for its effective use should therefore be intense (Endler 1992; 1993). This is particularly true for signal designs whose moment-to-moment appearance bears a heightened sensitivity to local conditions, such as structural colours (Kinoshita 2008; Mouchet and Vukusic 2018). Structural colours result from the selective reflectance of light by tissues organised at the nanoscale and can achieve levels of brightness and chromaticity that are unattainable with pigments alone (Kinoshita 2008; Johnsen 2012). Where coherent or quasi-coherent scattering is involved, as is most often the case, they are also capable of iridescence and limited-view expression (Prum 2006; Vukusic 2006).

Given that the optimal colour and/or brightness of iridescent signals is tied to the geometry of the signaller, viewer, and illuminant, its widespread use in nature suggests the existence of flexible behaviours for their effective presentation. That is, signallers should seek to capture available sunlight so as to maximise the conspicuousness of structurally coloured signals to viewers (Endler 1992; Endler and Thery 1996). This prediction has found broad support across model vertebrates with observational evidence showing how signallers vary the timing, location, or orientation of their displays to improve communication efficacy (Endler 1991; Endler and Thery 1996; Dakin and Montgomerie 2009; Bortolotti et al. 2011; Sicsu et al. 2013; Klomp et al. 2017; Simpson and McGraw 2018; O’Neill et al. 2019). Though this work is extremely valuable, a richer causal understanding awaits manipulative tests across temporal and taxonomic contexts.

Flies rank among the most diverse animal orders and this is reflected in the traits that have evolved in the service of visual communication (Marshall 2012). Sexually- and ecologically-specialised eye structures (Zeil 1983; Hardie 1986), elaborate colour patterns (Shevtsova et al. 2011; Butterworth et al. 2020), and ritualised display behaviours (Land 1993; Zimmer et al. 2003; Lunau et al. 2006; Butterworth et al. 2019) abound, and offer tractable but underutilised opportunities for general tests of theory. Work on visual signalling in the group continues to document the widespread presence of conspicuous structural colours, with recent efforts highlighting ‘wing interference patterns’ (WIPs) as potential vectors of information (Shevtsova et al. 2011; Katayama et al. 2014; Hawkes et al. 2019; Butterworth et al. 2020). These striking patterns are the result of thin film interference from chitin/air interfaces in the wing membrane (Shevtsova et al. 2011) and they are regularly presented during courtship (Land 1993; Frantsevich and Gorb 2006; Katayama et al. 2014; White et al. 2020). The wings bearing them are also typically transparent and transmit ca. 70-80% of incident light. This combination of iridescence and transparency makes their appearance uniquely environmentally contingent since bright visual backgrounds or the misalignment of signallers, receivers, and the sun will render the patterns invisible. The significance of visual backgrounds during signalling is well demonstrated (Katayama et al. 2014) and includes evidence for active choice in the wild (White et al. 2020). Whether and how signallers actively use the sun to improve communication, however, is unknown.

The genus *Lispe* is a cursorial group of muscid flies comprised of at least 163 species spanning most terrestrial biogeographic regions (Pont 2019). The group showcases striking interspecific and sexual variation in colouration and displays (Frantsevich and Gorb 2006; Pont 2019). *Lispe cana* is an exemplar endemic to the foreshores of Australia’s east coast. Males and females of the species bear structurally coloured UV-white and yellow faces, respectively, as well as dimorphic wing interference patterns (unpublished data). During their ground-based courtship male *Lispe cana* approach and straddle females from behind and maintain this position as females walk about the environment. They then release and move directly in front of females to present their faces and wings in a ritualised ‘dance’ (White et al. 2020). Males can and do enhance the salience of their signals through the selection of backgrounds against which to display, and similar potential benefits exist through the careful use of sunlight.

Here we use observational and manipulative field-based assays at a fine temporal scale to test the hypothesis that males should orient their displays toward the sun so to improve the salience (and, hence, attractiveness) of their iridescent signals. This predicts a bias in the orientation of male displays toward the sun’s azimuth position under clear skies, when the its position is readily visible and the potential benefits for signal salience are likely maximised. As a corollary, the natural or artificial removal of direct sunlight should disrupt any orientation bias by either impeding male’s ability to orient and/or diminishing the benefits of such behaviour for the salience of signals.

## Methods

### Sexual displays and orientations

Courtship in *Lispe cana* proceeds in four general stages: (1) males detect females when both parties are walking about their foreshore habitats; (2) males approach females from behind, before ‘straddling’ them and holding their wings closed as they continue to walk about the habitat; (3) males release females and rapidly move immediately in front of them at close distance (<5 mm), where they present their colourful wings and faces; (4) either the sequence returns to (2) with male re-straddling females, they mate, or (most often) they disperse. The second stage—straddling—thus offers males the opportunity to assess the environment and their orientation within it, while the third stage is when their decision to display at a given location and orientation is realised and is therefore the focus of our observations as detailed below.

We observed courtships in a population of *L. cana* that inhabits the supralittoral zone of Toowoon Bay, NSW, Australia (33.36 S, 151.50 E), in September and October of 2019. During each courtship, we recorded the orientation of the male during the display phase (described above; Fig. 1) of their ritualized dance, which we observed at a distance of 1-2 m. Courting pairs were haphazardly selected during slow, repeated walks by an observer along a 255 m transect at a distance of 5-10 m from the shoreline. Once a courtship naturally terminated with both parties dispersing, we placed a straight marker at the males’ final position in alignment with their medial body axis, before recording the markers’ orientation to the nearest one degree using a digital compass, as well as the time. To initially validate the accuracy of this approach we tested it against a video-based method in which male orientations were extracted from still-frames taken from 25 video-recorded courtships (with a GoPro Hero 6 at 30fps) in which a compass was visible in-frame. We found a strong correlation between orientations collected from video versus direct observation (Pearson’s r = 0.88), so we favoured the latter method for its convenience in garnering a large sample size.

**Figure 1:**
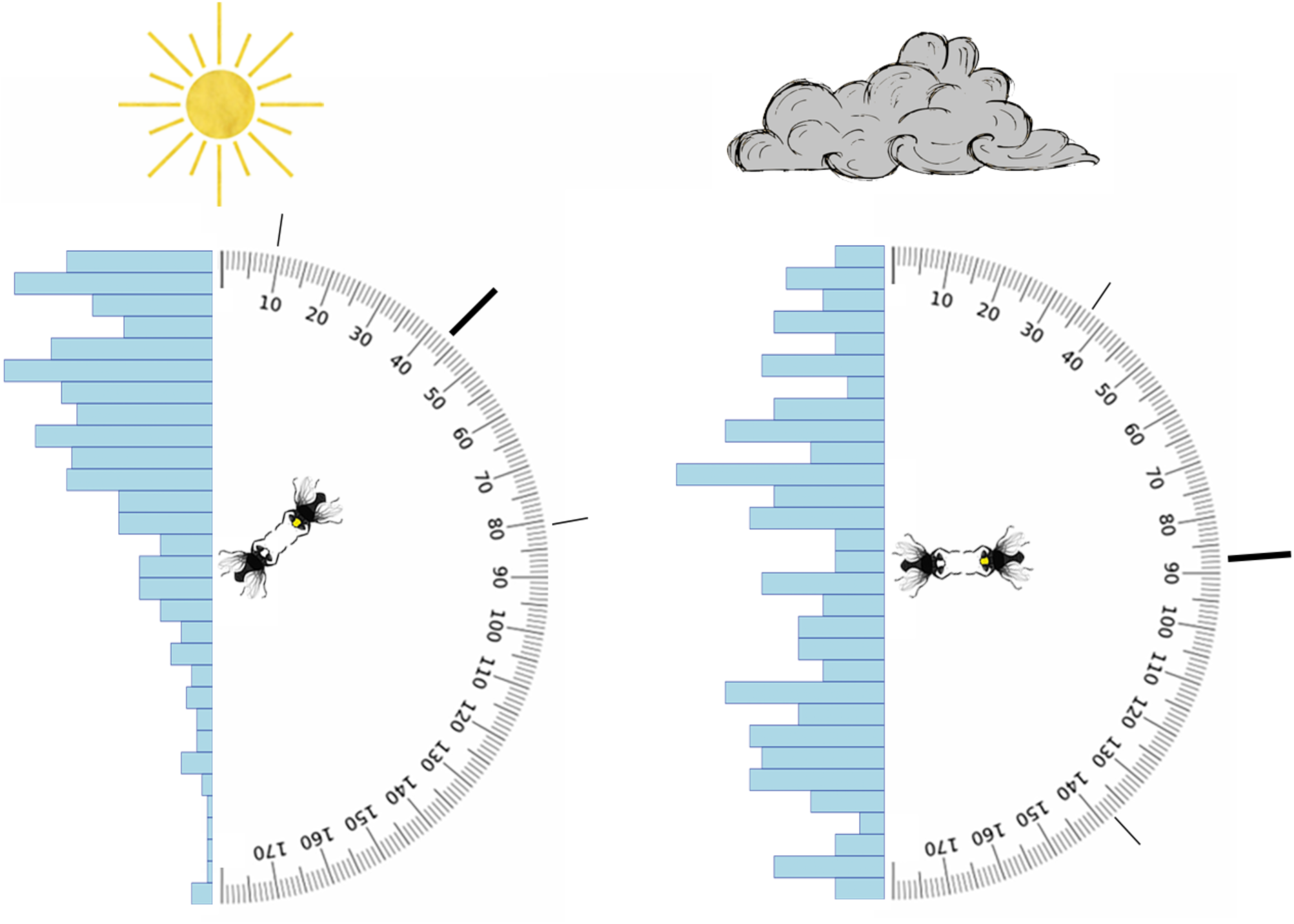
The orientations of *Lispe cana* during courtship displays under full sun (left) and heavy cloud (right). When displaying, male flies (white face) position themselves immediately in front of females (yellow face), and twist their wings forward in a seemingly ritualized ‘dance’. Angles denote the offset of the male midline from the solar azimuth during courtship, with 0° representing a display directly toward the sun, and 180° directly away. Histograms show the distribution of male offsets from the sun under both conditions (n = 450 full sun, n = 227 heavy cloud), with the median ± standard deviation indicated by solid lines.

**Figure 2:**
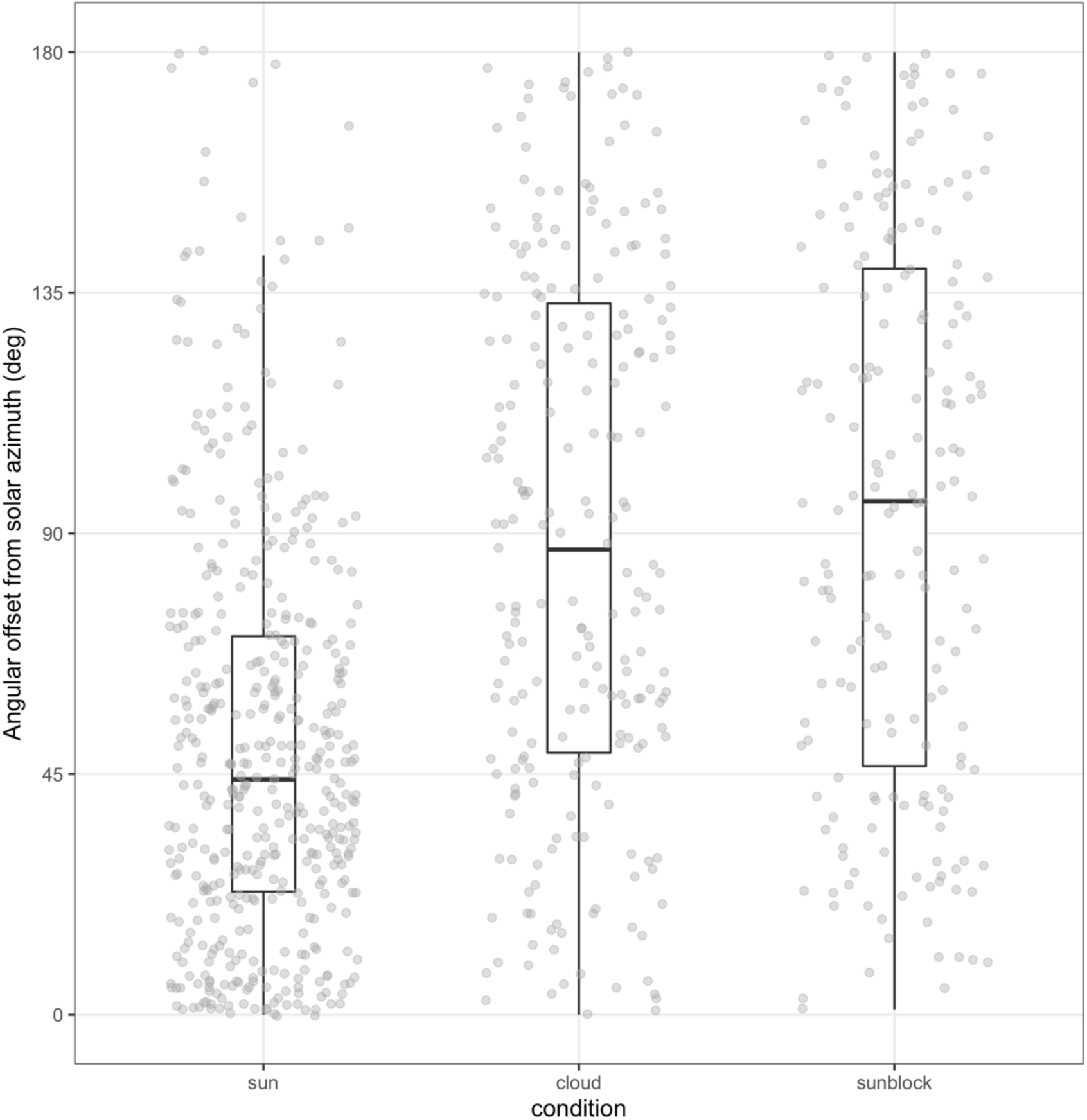
The orientation of displaying male *Lispe cana* with respect to the sun in conditions of full sun, heavy cloud, or experimentally obscured sunlight. Orientations are expressed as offsets from the solar azimuth, pooled across the 0730 - 1630 observation period, with 0° representing males presenting their iridescent signals directly toward the sun and 180° directly away.

Once courtships were observed, we used the time and location data to calculate the solar azimuth and elevation during every courtship via the package ‘suncalc’ (v. 0.5.0; Thieurmel and Elmarhraoui 2019) for R (v. 3.6.0; R Core Team 2018). We then converted our circular measure of male display orientations to a linear measure of the absolute deviation of each display from the sun’s azimuth. This gives a value between 0-180°, with 0° being a male displaying exactly toward sun, and 180° directly away from it. We ignore the position of females since they remain approximately opposite males at < 5 mm distance during displays (Fig 1).

To test the extent to which males can and do orient their displays we used the general procedure above across three conditions, both observational and manipulative. In the two observational conditions we recorded displays between the hours of 0730-1630 either on days of full direct sunlight (n = 451 observations across six days), or heavy cloud with the sun completely obscured (n = 227 across three days). For the manipulative assay we repeated the procedure as for full sun conditions but obscured any direct sunlight using a static black plastic sheet (Monarch pty. ltd) suspended on four aluminium poles ca. 10 m from the shoreline. This cast a ca. 3 × 3 m shadow throughout a given day, and we recorded only courtships that began and finished within this shaded area (n = 191 across three days). The broader environmental conditions were similar across days within treatments (summarised in supplementary table S1) and is partially accounted for in our statistical analyses (below).

### Solar orientation and signal luminance

To model the effects of male orientation on the salience of their signals we first recorded vector irradiances at the study site in 10° increments from 0° (directly toward the sun) to 180° (directly away) at 0800, under representative conditions of full sunlight and heavy cloud. We collected all spectra using an OceanInsight JAZ spectrophotometer with cosine-corrector and oriented the 180° collector perpendicular to the ground (i.e. in the same plane as male faces and wings during display). We then recorded the facial and whole-wing reflectance of 50 male *L. cana* using the JAZ with its pulsed-xenon light source, at an integration time of 40 ms and a boxcar width of 2. We used a bifurcated probe to both deliver and collect light, which we oriented normal to the face and wing planes at a working distances of ca. 5 mm. These spectra were lightly LOESS smoothed before being averaged to generate a representative male facial and whole-wing reflectance spectrum (Fig. S1).

We then estimated how the luminance of male wings and faces varies as a function of male orientation toward the sun. Here and throughout we focus on luminance alone given its importance over chromaticity (hue and saturation) for mating (White et al. 2020). To do so we calculated the integrated product of solar irradiance at each 10° increment from the solar azimuth, male facial or wing reflectance, and the Rh1-6 ‘achromatic’ receptor absorbance of *Musca domestica* (as the nearest analogue to *Lispe*). We conducted all spectral processing and visual modelling using lightr v1.1 (Gruson et al. 2019b) and pavo v2.4.0 (Maia et al. 2019) for R v3.6.1 (R Core Team 2018).

### Statistical analyses

To estimate whether male *Lispe cana* bias the orientation of their sexual displays toward the sun we used a generalized linear mixed-effect model fit by restricted maximum-likelihood, with Gaussian error and an identity link function. We specified male deviation from the solar azimuth (0-180) as the response which we square-root transformed. We did so after our inspection of errors from an untransformed model suggested slightly non-normal and heteroscedastic residuals, which were improved to within acceptable margins (as visually assessed) following transformation.

We specified the interaction between experimental condition (sun, cloud, and sun-blocked) and solar elevation as a fixed effect, along with the main effects of each. If males do actively orient their displays the result should be a negative effect of the full sunlight condition on male display offset (i.e. a tendency toward smaller offsets from the solar azimuth in full sun), when compared to the cloud and sun-blocked conditions. Our inclusion of solar elevation tests for reduced display accuracy at higher solar elevations when the sun’s azimuth is less readily discerned. Such an effect would only manifest under conditions of full sunlight when any display-orienting takes place, however, hence its inclusion via an interaction term. We specified the Julian date on which each measurement was taken as a random effect to account for variation associated with unmeasured within-day variables, such as broader climatic conditions.

To estimate the effects of male display orientation on signal luminance we first ran generalised additive models (GAM) fit by restricted maximum-likelihood, with the offset from solar azimuth (in 10° increments) as the lone predictor of signal luminance. The four resulting models describe the possible variation in male facial and wing luminance as a function of their sexual display orientation, during conditions of full sun and heavy cloud. We then used the results of the full-sun GAM alone to predict the luminance of male signals at their actual recorded orientations in both full-sun and heavy cloud conditions. We used only the full-sun GAM after finding no relationship between signal luminance and orientation under heavy cloud (Fig. 3d), and thus no possible benefits for signal salience to orienting toward the sun in such conditions (see results). We then used Kruskal-Wallis rank sum tests, with eta^2^ as an effect size, to estimate the difference in achieved luminance between male displaying under full sun and heavy cloud, with their luminance predicted from the full sun GAM. This, in effect, answers the question of what males achieve by orienting their displays under ideal conditions.

**Figure 3:**
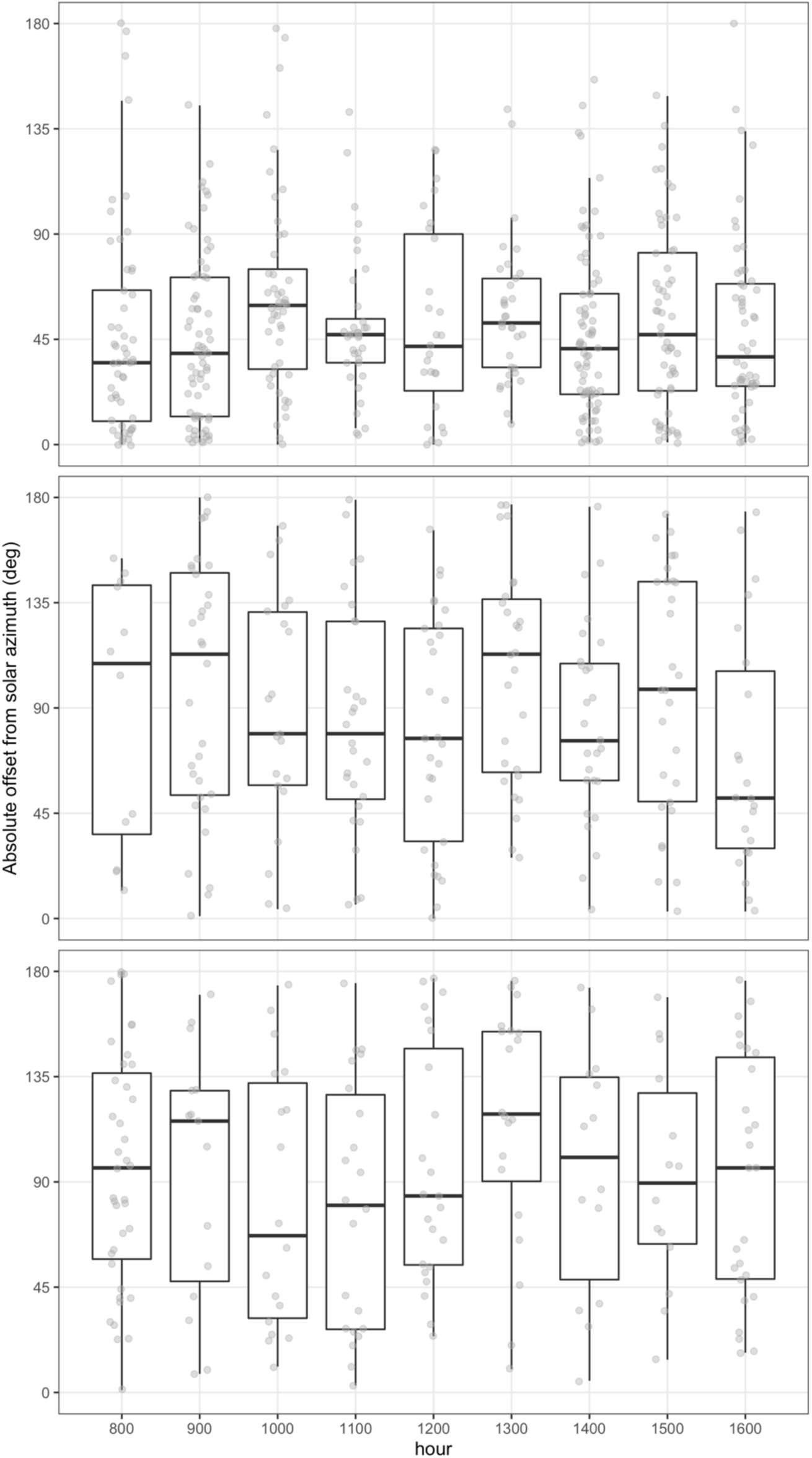
The orientation of displaying male *Lispe cana* in conditions of full sun (top), heavy cloud (middle), or experimentally obscured sunlight (bottom), pooled into hourly periods. Orientations are expressed as offsets from the solar azimuth, with 0° representing males presenting their iridescent signals directly toward the sun and 180° directly away.

### Data availability

All raw data and code are available via GitHub (https://github.com/EaSElab-18/ms_sunnyflies) and will be persistently archived upon acceptance.

## Results

We found clear differences in the orientation of male sexual displays across conditions (Fig. 1; Fig. 2). A likelihood-ratio test on the overall model identified a significant effect of experimental treatment alone (χ^2^ = 86.86, df = 2, *p* <0.001), with no evidence for an effect of solar elevation (χ^2^ = 2.54, df = 1, *p* = 0.111) nor any interaction between treatment and solar elevation (Fig. 3; χ^2^ = 0.81, df = 2, *p* = 0.665). Individual estimates (Table 1) show this was chiefly driven by a bias toward smaller angular offsets by displaying males under full-sun conditions alone (est = −3.11 ± 1.136, df = 9, t = −2.74, *p* = 0.02). Male displays were oriented toward the sun at a median offset of 44° ± 38 s.d. on days in which it was fully visible. But this effect was absent in conditions of heavy cloud (87° ± 50 s.d.) and artificially obstructed sunlight (96° ± 52 s.d.), which did not statistically differ from one another.

**Table 1:** Parameter estimates and summary statistics from a generalised linear mixed model examining the predictors of male display orientation in *Lispe cana.* Included are the experimental condition (full sun, heavy cloud, and artificially obscured sunlight) and solar elevation. Julian date was included as a random effect with a variance of 0.301. Full model conditional R^2^ = 0.169.

| Fixed effects | Estimate | SE | df | *t* | *p* |
| --- | --- | --- | --- | --- | --- |
| intercept | 8.389 | 0.771 | 853 | 10.88 | < 0.001 |
| condition (sun) | -3.115 | 1.336 | 9 | -2.743 | 0.023 |
| condition (sunblock) | 0.425 | 1.014 | 9 | 0.419 | 0.685 |
| solar elevation | 0.011 | 0.016 | 853 | 0.712 | 0.476 |
| condition (sun) x solar elevation | 0.014 | 0.023 | 853 | 0.624 | 0.533 |
| condition (sunblock) x solar elevation | -0.003 | 0.020 | 853 | -0.191 | 0.849 |

Our GAMs showed that the luminance of male wings (F_7.94_ = 356, p < 0.001) and faces (F_7.94_ = 355.50 p < 0.001) follow a near-identical sigmoidal relationship with solar orientation when illuminated in full sunlight (Fig. 4c). They are maximally luminant at 0° from the solar azimuth, inflect at ca. 45° (95% CI = 37-53° faces, 43-47° wings), before reaching their minima at ca. 80°. The luminance of wings, as compared to faces, varied across a slightly compressed range owing to their reduced reflectance (supplementary Fig. S1), but was otherwise strongly correlated with facial luminance. This effect was entirely absent in conditions of heavy cloud (Fig. 4d), as we found no relationship between either of facial (F1 = 0.190, p = 0.669) or wing (F1 = 0.192, p = 0.667) luminance and solar orientation.

**Figure 4:**
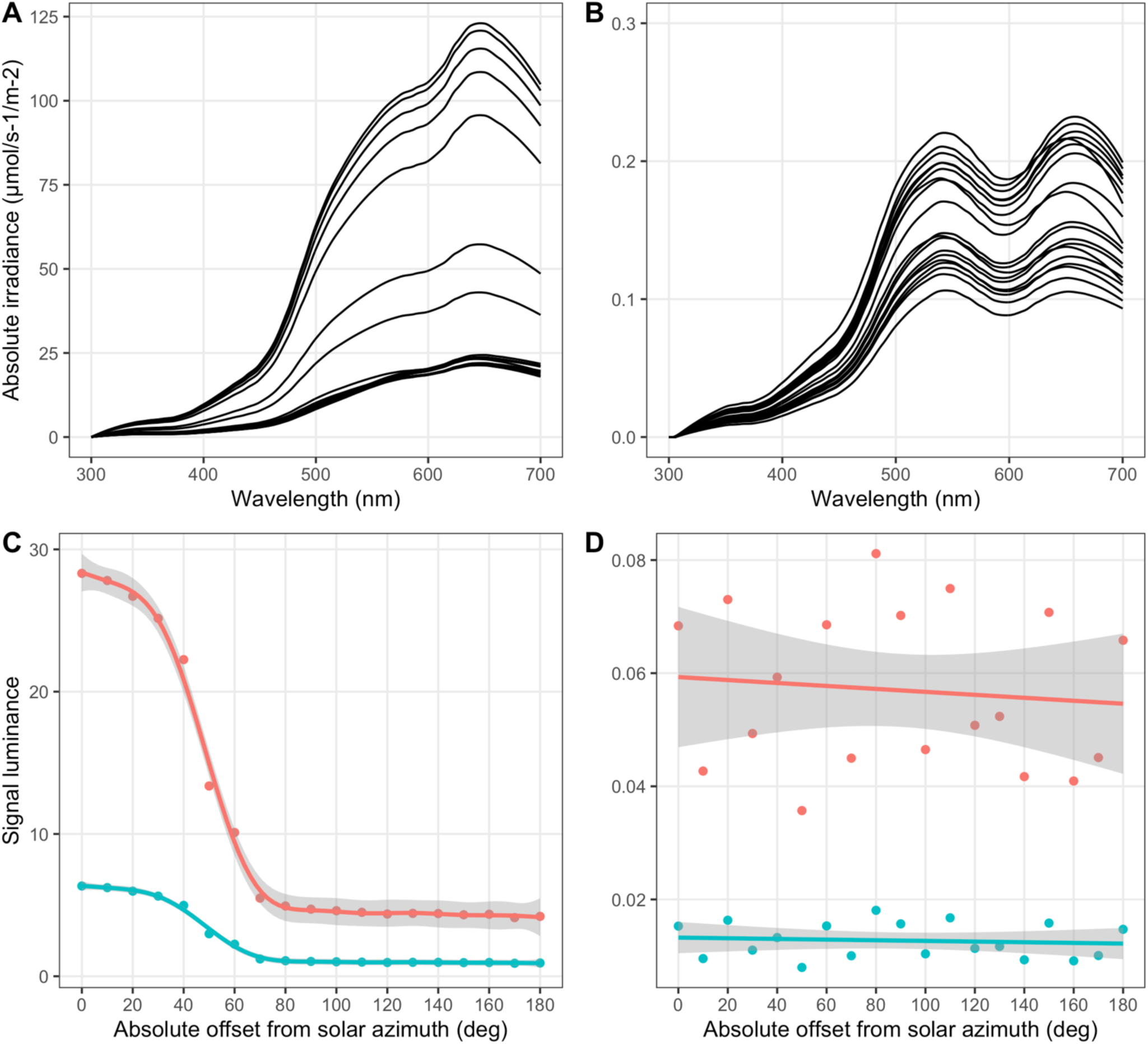
Solar irradiance and male signal luminance as a function of solar orientation. Solar irradiance in conditions of (a) full sun and (b) heavy cloud, recorded at 0° - 180° offset from the solar azimuth in 10° increments. Below is the luminance of male faces (red) and wings (green) across the same angular range, as modelled under full sun (c) or cloud (d). Lines denote generalised additive model fits ± standard errors. Not differing y axes across all panels.

Our subsequent test of signal luminance during actual male displays suggest clear benefits to active solar orientation in full sun. Both their faces (χ^2^ = 86.97, df = 1, p < 0.01, *η*^2^ = 0.127) and wings (χ^2^ = 86.70, df = 1, p < 0.01, *η*^2^ = 0.126) were subjectively brighter, with an approximately four-fold increase as a result of their active display orienting. Given the absence of a relationship between signal luminance and solar orientation under heavy cloud (as above), any active orientation by males could yield no benefits to signal salience in such conditions.

## Discussion

The intensity and composition of sunlight shapes the salience of visual signals, and selection should favour its flexible use. Here we tested this prediction in the cursorial fly *Lispe cana* whose structurally coloured faces and wings are central to their ritualised courtship displays. Our field-based observations show that males bias the orientation of their signals toward the sun under clear skies, with individuals displaying at approximately 44° offset from the solar azimuth on average. This degree of precision is maintained throughout the day irrespective of solar elevation. Notably, however, the directional bias in displays broke down in conditions of heavy cloud, and we were able to reconstruct this effect by experimentally obstructing the sun under otherwise-ideal conditions. Our visual modelling of signal luminance identified clear benefits to male display behaviour, with an approximately four-fold difference in the fly-subjective luminance of male signals across the 180° range of possible orientations. These combined results offer causal evidence that male flies actively orient their conspicuous displays toward the sun, with tangible consequences for the salience of their structurally coloured signals. Below we consider our findings in turn and discuss their relevance for communication efficacy and signal evolution more broadly.

One immediate question is how male *L. cana* reliably orient their displays throughout the entire day. The sun is the brightest spot in the sky so tracking its position directly is an obvious solution, and this information can be supplemented through the use of indirect cues such as intensity, chromaticity, and polarization gradients. On clear days, the intensity of sunlight increases predictably toward the solar azimuth and atmospheric scattering means the sky is relatively richer in long-wavelength light in the solar, as opposed to anti-solar, hemisphere (Fig. 3a; Coemans et al. 1994). The degree of polarisation is similarly graded and increases with the angular distance from the sun (Coemans et al. 1994). The integrated use of such cues is well described among insects, including flies (Philipsborn et al. 1990; Weir and Dickinson 2012), albeit typically in the context of navigation. Monarch butterflies, for example, rely on direct and indirect skylight cues to calibrate their time-compensated sun compass during annual migrations (Heinze and Reppert 2011; Reppert et al. 2016), while desert ants integrate solar intensity, chromaticity, and polarisation during navigation to and from the nest (Müller and Wehner 2007; Lebhardt and Ronacher 2015).

Though our results cannot directly test whether male *L. cana* make use of all available cues when orienting, the suppression of their orientation bias with the obstruction of direct sunlight—both artificial and natural—suggests that they first and foremost rely on the sun’s immediate position. This follows from the fact that gradients in each of polarization, and (to a lesser extent) luminance and chromaticity are present even when the sun is not directly visible, and many flies are able to use this information to guide behaviour in other contexts (Philipsborn et al. 1990; Weir and Dickinson 2012). However, while there is reason to expect that males *should* be able to orient their displays in non-optimal conditions, our modelling suggests that there is no benefit in doing so. Signal luminance bears no relationship to solar orientation under heavy cloud (Fig. 3d), so males should instead prioritise more consequential effects such as the selection of suitable display backgrounds (White et al. 2020). Whether they do so remains to be tested.

The importance of accurate solar orientation during courtship is heightened by *L. cana*’s use of structurally coloured ornaments. This is particularly true for wing interference patterns, the colour and luminance of which is a product of thin-film interference from the layering of chitinous wing membranes (Shevtsova et al. 2011). Since these colours are highly specular (i.e. dominated by mirror-like reflection) in nature, the absence of a point-source illuminant such as the sun can render the patterns indistinguishable (Shevtsova et al. 2011). Males that are less able to optimise their orientation would therefore suffer penalty not only in the quality of their wing colouration (e.g. via reduced intensity), but the ‘category’ too, as the patterns will be entirely absent at oblique illumination angles.

Since males’ wings are also semi-transparent, their optimal appearance is equally dependent on the background against which they are presented. Dark backgrounds—such as seaweed common to the foreshore habitats of *L. cana*—are fit for purpose, as they minimise transmitted light which would overwhelm the weakly reflective interference patterns. Recent work has documented just such an effect with males preferentially displaying against darker visual backgrounds, and the magnitude of signal/background contrast predicting mating success (White et al. 2020). Though we did not track mating success here a comparable effect on fitness is plausible, given that imprecision in either background selection or orientation relative to the sun will produce a similarly diminished signal (Fig. 4c). It remains to be seen whether and to what extent the sun’s position and visual backgrounds are weighed and integrated during courtship, but the accumulating evidence paints a striking picture of behavioural flexibility in service of communication efficacy.

The behaviours dedicated to the presentation of iridescent faces and wings imply an important role for the latter in sexual communication, though their precise function is as-yet unresolved. As structurally coloured ornaments the macro-scale appearance of faces and WIP’s is closely tied to the nanoscale organisation of underlying structures (Kinoshita 2008). If males differentially vary in their ability to secure the conditions and/or material required to generate such highly organised tissues, then the expression of facial and wing signals may be condition dependent (Zahavi 1975; Keyser and Hill 1999; Shawkey et al. 2003).

In *Lispe cana* the male’s UV-white are a product of incoherent scattering by modified bristles (unpublished data, see Frantsevich and Gorb 2006 for details in a sister species), the brightness of which will largely depend on the size and density of scatterers (Giraldo and Stavenga 2007; White et al. 2012; Stuart-Fox et al. 2018). The appearance of their wing interference patterns is instead a product of coherent scattering, as noted above, with colour differences across the wing surface chiefly determined by variation in the thickness of the cuticular thin-films (Shevtsova et al. 2011). Both are constructed and fixed during ontogeny using the pool of resources gathered during the larval stage, and so their final appearance may reflect individual variation in foraging ability and condition more generally (e.g. Kemp 2008; Kemp and Rutowski 2007). Recent meta-analytic evidence supports the potential for condition-dependent expression among structural colours in general (White 2020), and also suggests that the greater organisation demanded by coherent, as opposed to incoherent, scatterers makes them particularly suitable as condition-dependent ornaments. This argues for WIPs as more probable vectors of information on mate quality in *L. cana* than faces, with the latter potentially serving as signals of sex or species identity. Despite convincing evidence for sexual selection in WIP evolution (Katayama et al. 2014; Hawkes et al. 2019), however, direct tests of their function as signals remain outstanding.

In addition to iridescent ornaments as conduits of information, a non-exclusive possibility is that male behaviour is itself informative of mate quality. Courtship displays are likely to be energetically costly as the cycle of pursuing, straddling, and displaying with a given female is often repeated several times before a clear outcome is achieved. Males clearly vary in their ability achieve and sustain optimal orientations during courtship (Fig 2). If this variation is dependent on male condition or ‘quality’ then their coloured ornaments may simply serve as amplifiers of their behavioural performance. Displays against sub-optimal backgrounds or at imprecise solar orientations would be readily apparent, particularly in the appearance of WIPs (Shevtsova et al. 2011). Potential benefits to selective females are predictable since, as predators, flight ability is central to prey capture in *L. cana* (Steidle et al. 1995; Pont 2019). This form of condition-dependent variation in courtship effort, and corollary benefits to choosy viewers, is well documented (Kotiaho 2000; Wagner and Hoback 1999; Jennions 1998). Similar effects are known for variation in the precision of displays, such as in the quality of courts constructed by bowerbirds (Endler et al. 2010; Kelley and Endler 2012), and the combination of limited-view iridescence and precision displays in *L. cana* creates potential for a similar dynamic.

Our results extend a growing body of evidence which suggests extensive coevolution between signals and display behaviours (Endler 1991; Endler and Thery 1996; Dakin and Montgomerie 2009; Bortolotti et al. 2011; Sicsu et al. 2013; Klomp et al. 2017; White 2017; Simpson and McGraw 2018). Notably, the strength of this link appears to vary with the extent of fluidity in signalling environments and/or signals. That is, signals and environments which vary dynamically across fine temporal or spatial scales are frequently coupled with flexible behaviours to enhance the efficacy of information exchange. Iridescent signalling systems—such as that examined here—fall at one extreme, with examples from insects (White et al. 2015), birds (Endler and Thery 1996; Dakin and Montgomerie 2009; Simpson and McGraw 2018), and guppies (Endler 1991) revealing considerable plasticity in display behaviour for optimising signal delivery. Environments which vary unpredictably across fine spatial and temporal scales appear to favour similar adaptive solutions. The foreshores of *L. cana* offer a case in point, since a single wave can add or remove tracts of seaweed which the flies use for shelter and signalling, and thus entirely restructure their immediate visual environment.

Despite these few well-defined examples, the extent to which we can we can predict any one of behaviour, habitat structure, or signal design from knowledge of the others is a long-standing question (Poulton 1890; Lythgoe 1988; Endler and Thery 1996). One key to resolving this challenge is a deeper appreciation of the spatio-temporal complexity of visual communication. Exciting advances continue to be made in defining and integrating the spectral (Maia et al. 2019; van den Berg et al. 2019), spatial (Caves et al. 2018; Stoddard and Osorio 2019), and temporal (Gruson et al. 2019a) features of signal production and perception, and systems such as *Lispe cana* offer a promising context for empirical progress.

## Acknowledgments

TEW is grateful to Elizabeth Mulvenna and Cormac White for their support.

