## Supplementary material for "Flies improve the salience of iridescent sexual signals by orienting toward the sun"

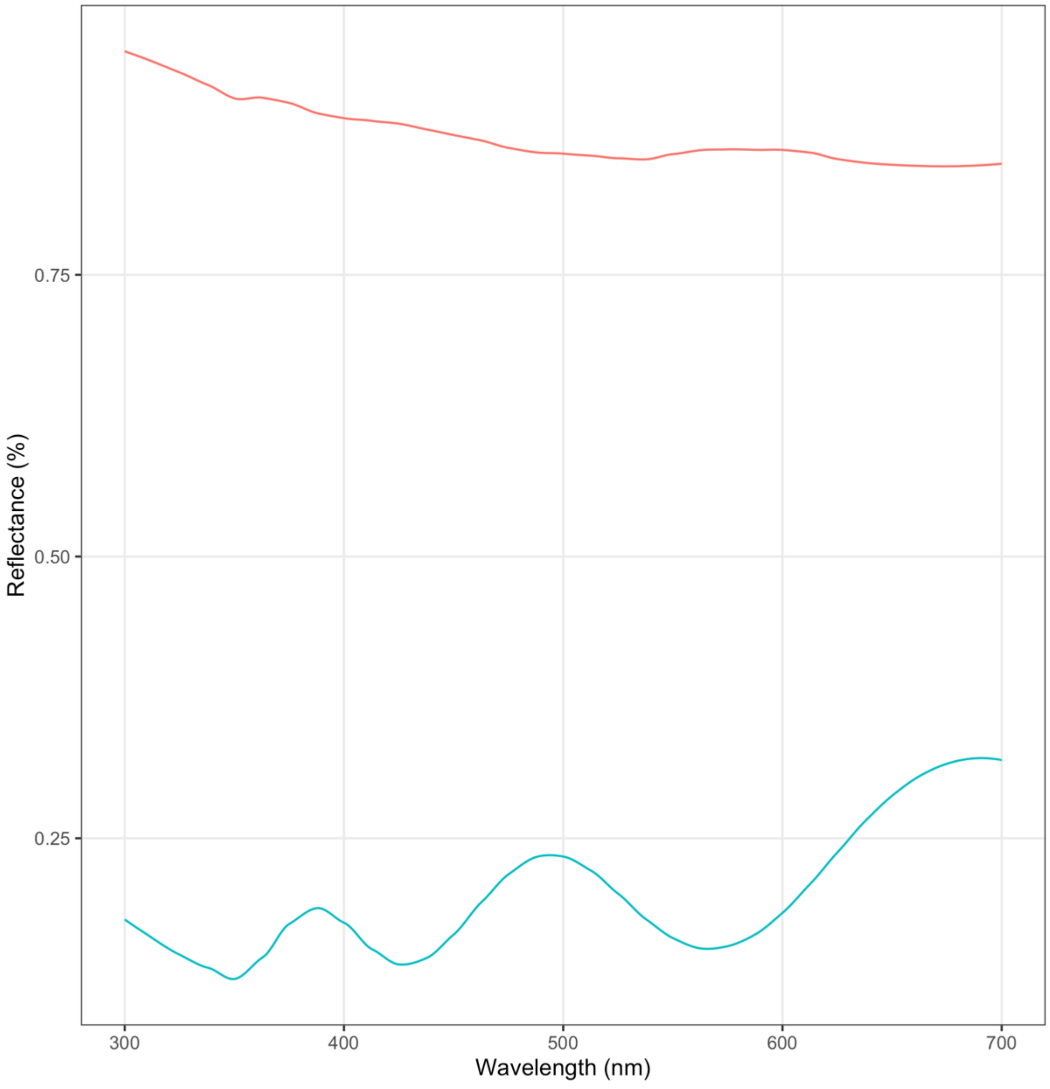


**Figure S1:** Normalised reflectance spectra of the structurally coloured faces (red) and wings (blue) of male *Lispe cana.*

**Table 1:** Diel summary environmental data during each condition. Measures are drawn from the Australian Bureau of Meteorology's Wyong Bridge Upstream station, situation 9.8 km from the study site (accessible at http://www.bom.gov.au/climate/data/).

| **Condition** | **Temperature (°C)** | | | | **Solar exposure (MJ/m$^2$)** | | | | **Rainfall (mm)** | | | |
|---|---|---|---|---|---|---|---|---|---|---|---|---|
| | min | max | mean | sd | min. | max | mean | sd | min. | max | mean | sd |
| sun | 19.8 | 33.3 | 27.3 | 5.2 | 18.9 | 25.4 | 23.0 | 4.4 | 0 | 0 | 0 | 0 |
| cloud | 17.0 | 23.3 | 19.5 | 2.7 | 10.9 | 21.4 | 15.2 | 1.8 | 0 | 0 | 0 | 0 |
| sun-block | 32.7 | 34.3 | 33.2 | 0.77 | 25.2 | 26.6 | 26.0 | 0.6 | 0 | 0 | 0 | 0 |
